# Identification of positive chemotaxis in the protozoan pathogen *Trypanosoma brucei*

**DOI:** 10.1101/667378

**Authors:** Stephanie F. DeMarco, Edwin A. Saada, Miguel A. Lopez, Kent L. Hill

**Affiliations:** Molecular Biology Institute, University of California, Los Angeles, California, USA; Department of Microbiology, Immunology and Molecular Genetics, University of California, Los Angeles, California 90095, USA; California NanoSystems Institute, University of California, Los Angeles, California, USA

## Abstract

To complete its infectious cycle, the protozoan parasite, *Trypanosoma brucei*, must navigate through diverse tissue environments in both its tsetse fly and mammalian hosts. This is hypothesized to be driven by yet unidentified chemotactic cues. Prior work has shown that parasites engaging in social motility *in vitro* alter their trajectory to avoid other groups of parasites, an example of negative chemotaxis. However, movement of *T. brucei* toward a stimulus, positive chemotaxis, has so far not been reported. Here we show that upon encountering *E. coli,* socially behaving *T. brucei* parasites exhibit positive chemotaxis, redirecting group movement toward the neighboring bacterial colony. This response occurs at a distance from the bacteria and involves active changes in parasite motility. By developing a quantitative chemotaxis assay, we show that the attractant is a soluble, diffusible signal dependent on actively growing *E. coli*. Time-lapse and live video microscopy revealed that *T. brucei* chemotaxis involves changes in both group and single cell motility. Groups of parasites change direction of group movement and accelerate as they approach the source of attractant, and this correlates with increasingly constrained movement of individual cells within the group. Identification of positive chemotaxis in *T. brucei* opens new opportunities to study mechanisms of chemotaxis in these medically and economically important pathogens. This will lead to deeper insights into how these parasites interact with and navigate through their host environments.

**Importance:** Almost all living things need to be able to move, whether it is toward desirable environments or away from danger. For vector-borne parasites, successful transmission and infection require that these organisms be able to sense where they are and use signals from their environment to direct where they go next, a process known as chemotaxis. Here we show that *Trypanosoma brucei*, the deadly protozoan parasite that causes African sleeping sickness, can sense and move toward an attractive cue. To our knowledge, this is the first report of positive chemotaxis in these organisms. In addition to describing a new behavior in *T. brucei*, our findings enable future studies of how chemotaxis works in these pathogens, which will lead to deeper understanding of how they move through their hosts and may lead to new therapeutic or transmission-blocking strategies.

## Introduction

A fundamental aspect of virtually all motile organisms is the ability to move in response to a change in the environment. One strategy to do this is through chemotaxis, the movement of an organism toward or away from a chemical cue. In microbial systems, chemotaxis has been best characterized in bacteria and social amoeba, which both employ chemotaxis to locate nutrients and avoid unfavorable environments (1–3). Many bacterial pathogens in particular rely on chemotaxis to move toward their desired site of infection (4–7). For protozoan pathogens, which typically must navigate through multiple hosts and a variety of different tissues in each host, chemotaxis has also been hypothesized to be necessary for pathogenesis and transmission (8–13).

*Trypanosoma brucei* is a protozoan pathogen that causes African sleeping sickness in humans and nagana in cattle. *T. brucei* is transmitted to a mammalian host through the bite of an infected tsetse fly. In the mammalian host, the parasite first mounts a bloodstream infection before penetrating the blood vessel endothelium to enter the central nervous system, resulting in lethality if not treated (14). *T. brucei* also infiltrates adipose and dermal tissue and these extravascular sites represent biologically significant parasite reservoirs that may influence pathogenesis and transmission (15–17). Within the tsetse fly vector, *T. brucei* must complete an ordered series of directional migrations through specific host tissues in order to be transmitted to a new mammalian host (18). Mechanisms underlying tissue tropisms observed in the mammalian host and insect vector are unknown.

Evidence demonstrating that *T. brucei* can adjust its motility in response to external cues comes from *in vitro* studies of social motility (SoMo), which occurs in procyclic-form *T. brucei* (tsetse fly midgut stage) when cultivated on semi-solid agarose (19). During SoMo, *T. brucei* cells assemble into groups that engage in collective motility, moving outward from the point of inoculation to form radial projections. Movement outward is cell density-dependent, suggesting a quorum sensing component to control of motility (20). Furthermore, when parasites in projections sense other *T. brucei* cells, they actively avoid one another, either by stopping their forward movement or changing their direction of movement, thus exhibiting capacity for negative chemotaxis (19, 21). Additional work revealed that SoMo depends on cAMP signaling in the flagellum (22–24), and recent *in vivo* work has demonstrated that flagellar cAMP signaling is required for *T. brucei* progression through fly tissues (8). Thus, simply being able to move is not sufficient to complete the transmission cycle and the combined findings support the idea that *T. brucei* depends on chemotaxis in response to extracellular signals to direct movement through host tissues. To our knowledge however, positive chemotaxis has not been reported for *T. brucei*.

Here we report that *T. brucei* engaging in SoMo exhibit positive chemotaxis toward *E. coli*, a behavior we term “BacSoMo.” While *T. brucei* does not typically interact with *E. coli* in its natural hosts, it does encounter other bacteria, and *E. coli* serves as an easy-to-control bacterial sample for use in dissecting chemotaxis *in vitro*. We find that the response is mediated by an active change in parasite motility that occurs at a large distance from the bacteria, indicating response to a chemical cue. Supporting this idea, we show that attraction is mediated by a signal that diffuses through the culture medium and requires actively growing *E. coli*. Our findings allowed us to begin dissecting cellular behavior that underlies chemotaxis in *T. brucei*, revealing changes in motility at both the group and individual cell level. We expect these studies to lead to a deeper understanding of how trypanosomes navigate through diverse environments encountered during their transmission and infection cycle.

## Results

### 1. Socially behaving *T. brucei* exhibit chemotaxis toward *E. coli*

During social motility (SoMo), *T. brucei* cells engage in collective motility to form radial projections that have a clockwise curvature (when viewed from above, Fig. 1A, left) (19). When parasites in these projections sense other *T. brucei* cells, they actively avoid one another, either by stopping their forward movement, or by changing their direction of movement (Fig. 1A, center) (19). We found, however, that when encountering *E. coli* on the SoMo plate, the parasite projections continue moving to make contact with the bacteria, and even appear to alter their movement to move directly toward the bacteria (Fig. 1A, right).

**Figure 1:**
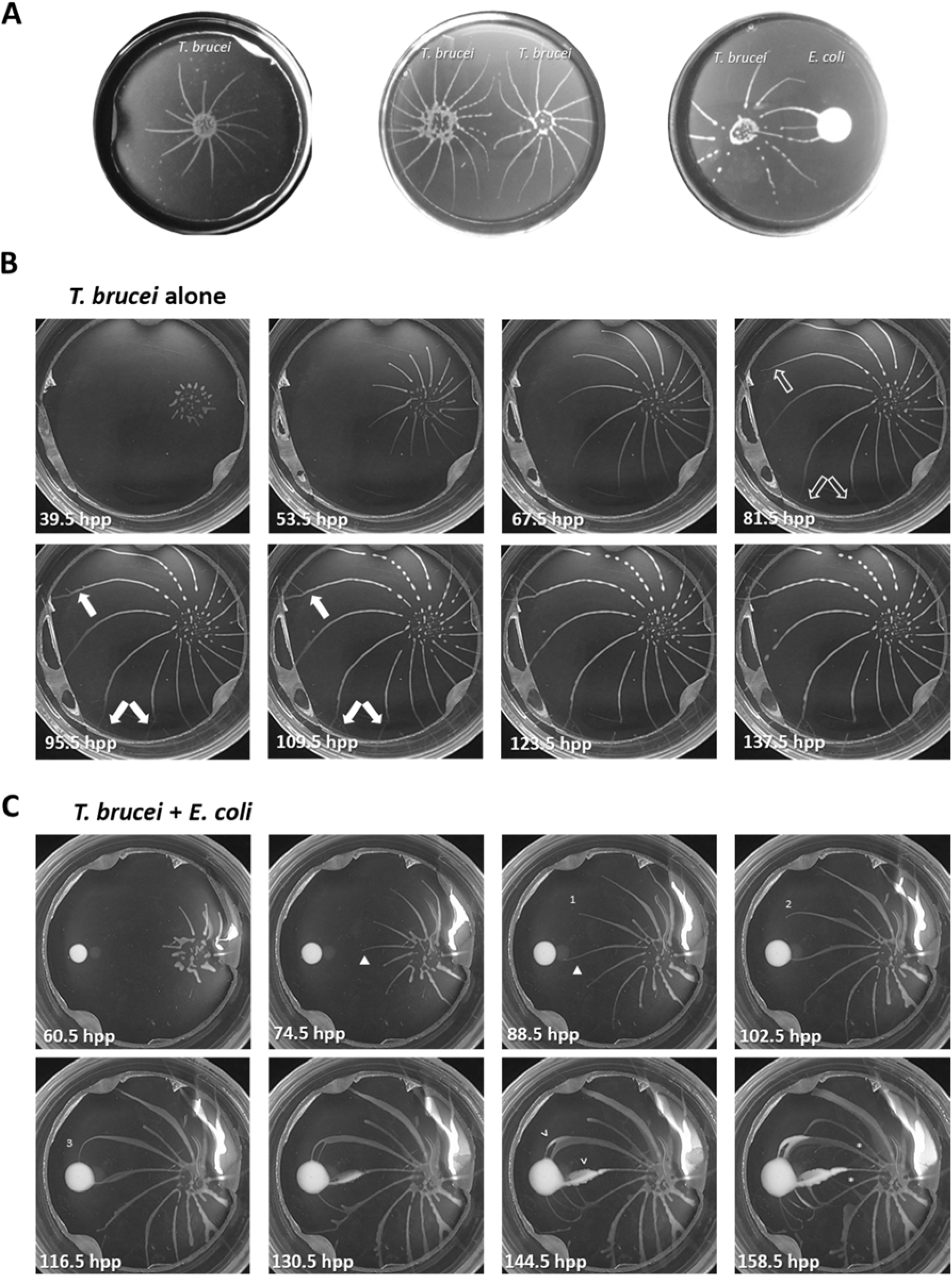
Socially behaving *T. brucei* is attracted to *E. coli*. A) *T. brucei* on a semi-solid surface engages in social motility (SoMo) (left). Projections of two groups of *T. brucei* originating from the same suspension culture are repelled by one another (center). *T. brucei* is attracted to *E. coli* (right). B) Stills from a time-lapse video of *T. brucei* engaging in SoMo (Movie 1). Unfilled arrows point to projections before branching. Filled arrows point to projections that have formed branches. C) Stills from a time-lapse video of *T. brucei* exhibiting positive chemotaxis toward *E. coli* (Movie 2). Time-stamps are indicated in hours post-plating (hpp). Numbers 1-3 indicate a projection that alters its path in response to *E. coli*. Closed arrowhead points to a change in curvature of the projection as it changes its path. Open arrowheads point to locations where *E. coli* has entered the projections. Asterisks indicate regions where a projection has crossed a different projection.

We hypothesized that the movement toward bacteria is chemotactic in nature, but we also considered whether it might instead reflect preferential growth in the direction of bacteria. To distinguish between these possibilities, we used time-lapse imaging to examine dynamics of *T. brucei* movement. SoMo assays were performed with or without bacteria. Images were taken every 30 minutes over the course of 162 hours (Fig. 1B, Movie 1) or 185 hours (Fig. 1C, Movie 2) and compiled into movies. Plates were turned upside-down to prevent condensation on the lid from interfering with the analysis, and then imaged from the bottom side of the plate. Therefore, the curvature of the projections observed in videos and time-lapse images is counter-clockwise.

In the absence of bacteria, parasites moved continually outward, forming arced projections that radiated away from the inoculation site and rarely altered their general direction of movement (Fig. 1B and Movie 1). At early stages, projections maintained relatively even spacing and uniform width, having a single leading edge without branching. As projections neared the periphery, the space between projections increased and parasites advanced from the lateral edge to form branches (arrows Fig. 1B; Movies 1 and 3). The observation that branching only occurred when spacing between neighboring projections increased supports the idea (19) that inhibitory signals from parasites in adjacent projections prevent parasite movement from the side of projections. In some cases, thickening of a projection was observed prior to branching (Fig. S1, Movie 3), suggesting that cell density-dependent signals driving parasite movement outward (20) may overcome inhibitory signals between projections. Parasites in branches continued to adjust their movements so that they did not make contact with adjacent parasites (Fig. 1B, Fig. S1, Movies 1 and 3), demonstrating that parasite-dependent inhibitory signals were still active.

Time-lapse imaging of SoMo assays carried out in the presence of a neighboring bacterial colony allowed us to define the point at which parasites sense and respond to bacteria, both spatially and temporally, and this revealed several important findings. First, these analyses clearly demonstrate that parasite movement toward bacteria is an active response and not simply an absence of avoidance. Notice for example, that rather than continuing to the periphery of the plate, as occurs in the absence of bacteria, parasite projections curve sharply to move directly toward bacteria (Fig. 1C: Numbers 1-3, 88.5 – 116.5 hpp). Moreover, in the presence of bacteria, projections change curvature from counterclockwise to clockwise (Fig. 1C: triangle, 74.5 and 88.5 hpp) and exhibit extensive branching (Fig. 1C, 130.5 hpp – 158.5 hpp), with branches moving directly toward the bacteria. Second, these changes in movement occur at a large distance from the edge of the bacterial colony, indicating that parasites are responding to a chemical cue derived from the bacteria, rather than detecting the bacteria by direct contact (Fig. 1C, Movie 2).

A third important result to come from time-lapse studies is that the timescale of the response rules out the possibility that movement toward bacteria simply represents preferential growth in this direction. For example, between 88.5 and 116.5 hpp (Fig. 1C: Numbers 1-3), parasites have turned sharply toward bacteria and advanced to contact the bacterial colony. This projection impacted the bacteria at 109 hpp (Movie 2). Growing on plates, *T. brucei* has a doubling time of approximately 24 hours (19), thus the 20.5-hour time interval between 88.5 hpp and 109 hpp, thus represents slightly less than one cell doubling time, yet the parasites moved 22.61 mm (Movie 2), which corresponds to approximately 1046 cell lengths (25). Clearly this distance cannot be accounted for by cell doubling. Therefore, parasites alter their movement in response to a signal that acts at a distance to move directly toward bacteria.

In the moments before parasite projections impact the bacterial colony, we see individual parasites move directly from parasite projections into the bacterial colony (Movie 5, part 1). After contact, parasites spread out as they infiltrate the bacterial colony (Movie 5, part 2). Meanwhile, bacteria from the colony advance outward along parasite projections (Fig. 1C: arrowheads). As bacteria advance along one projection, parasites from adjacent projections become attracted to the position now occupied by bacteria (Fig. 1C, 144.5 and 158.5 hpp). Therefore, repositioning of the bacterial population directly correlates with a change in position of the attractant source. As parasites move to this attractant, they now even cross other projections of parasites to reach the bacteria (Fig. 1C: asterisks, 158.5 hpp), a phenomenon never observed in the absence of attractant. This indicates that the attractive cue from the bacteria is stronger than the repulsive cue that otherwise prevents contact and crossing of projections (19). Altogether, time-lapse video analysis demonstrated that parasites in projections are not exhibiting preferential growth towards *E. coli*, but are actively directing their movement toward it through positive chemotaxis in response to an attractant that acts at a large distance from the source.

### 2. The attractant is diffusible and requires actively growing E. coli

To define characteristics of the attractant, we employed a quantitative chemotaxis assay (Fig. 2A) developed based on similar assays used to study chemotaxis in parasitic worms (26). In this assay a chemotactic index is calculated for each sample by determining the number of projections that enter a 2-cm diameter centered around the sample, compared to how many projections enter a circle centered at the same position when no sample is present. A positive chemotaxis index indicates attraction, while a negative index indicates repulsion, and “perfect” attraction or repulsion is defined as + 1 or – 1, respectively. In this assay, a colony of live bacteria has a positive chemotactic index, + 0.37, indicating *T. brucei* is strongly attracted to *E. coli*. We previously showed that *T. brucei* is repelled by other groups of *T. brucei* (19). Therefore, as a negative control, we used a *T. brucei* PDEB1 knockout mutant that does not form projections (8) and produces a colony approximately equal in size to bacterial colonies grown on SoMo plates (Fig. 2A). We found that *T. brucei* was “perfectly” repelled by PDEB1 KO *T. brucei*, with a chemotaxis index of – 1, supporting the capacity of the assay to distinguish attractive versus repulsive chemotaxis.

**Figure 2:**
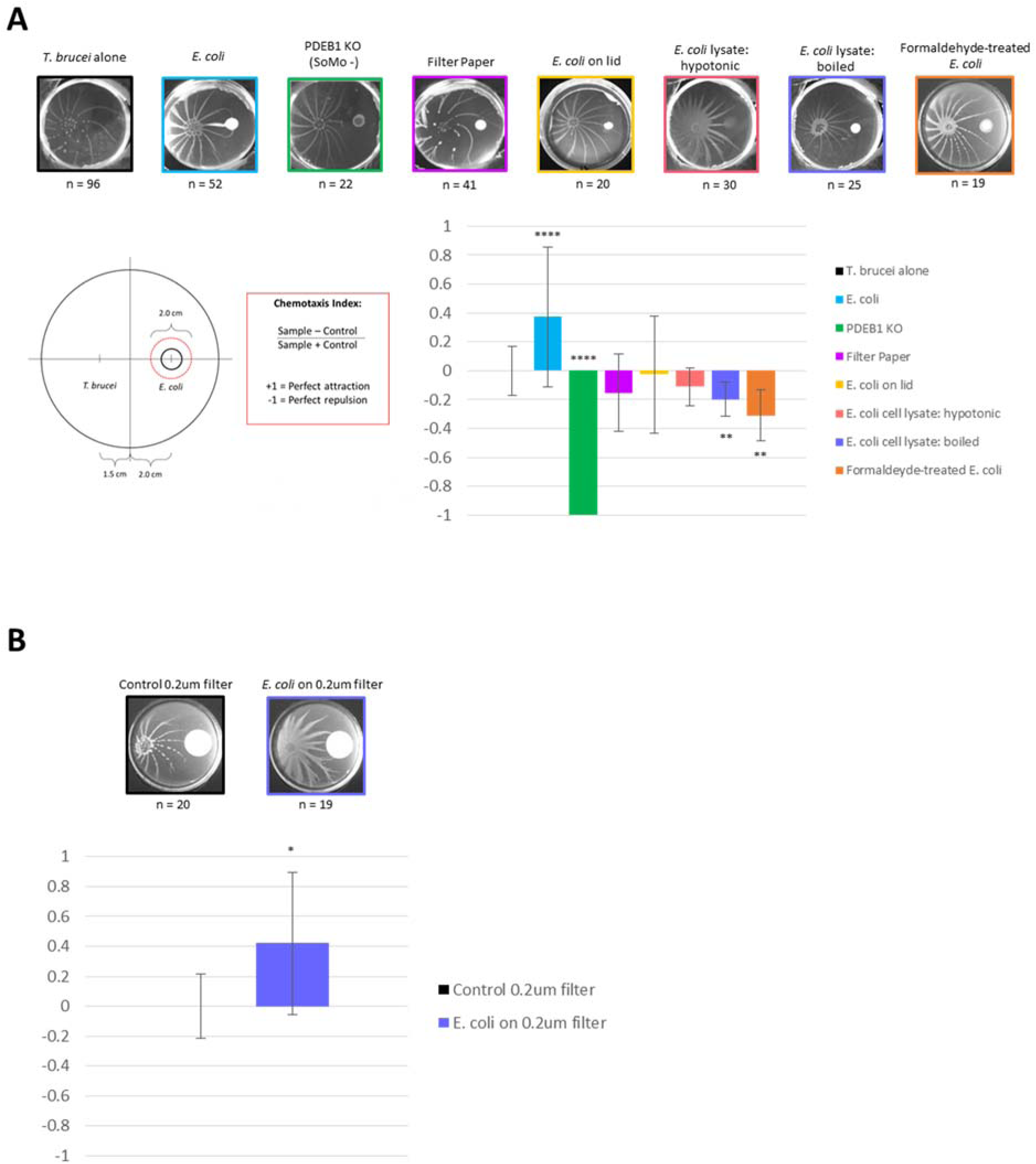
The attractant is diffusible and requires actively growing *E. coli*. A) Requirements for attraction were quantified by a chemotaxis index (diagram at lower-left), defined as the number of projections entering the 2-cm diameter red circle in the experimental sample subtracted by the number of projections entering the same circle on control plates (*T. brucei* alone), divided by the total number of projections in both samples. Representative pictures of each condition tested are shown (upper row) with their chemotaxis indices (lower-right). Error bars represent SEM. Unpaired two-tailed t-test with Welch’s correction was used to measure significance compared to the *T. brucei* alone control: **** p < 0.0001, ** p < 0.01. B) A chemotaxis index was calculated for *T. brucei* in response to *E. coli* growing on a 0.2µm filter compared to a 0.2µm filter alone. Error bars represent SEM. Unpaired two-tailed t-test with Welch’s correction was used to measure significance compared to the *T. brucei* in response to 0.2µm filter alone control: * p < 0.05.

We next considered whether attraction to bacterial colonies might simply reflect a response to a physical perturbation in the agarose surface created by the physical presence of the bacterial colony. To assess this, the chemotactic index of a piece of filter paper was tested, and *T. brucei* showed no significant chemotactic response (Fig. 2A), reinforcing the hypothesis that the response to bacteria is a chemotactic response.

Bacteria can produce both volatile – released into the air – and soluble compounds, which can serve as chemotactic cues (27, 28). To differentiate between these, we first asked whether *T. brucei* would respond positively to aerosolized volatile compounds. To do this, we employed a variation in the chemotaxis assay in which *E. coli* was plated on the lid of the petri dish while *T. brucei* was inoculated on the bottom. *T. brucei* showed no response to *E. coli* on the lid, indicating that the attractive cue is non-volatile and suggesting it is a soluble factor that diffuses through the culture medium.

To test if the attractant was diffusible through the culture medium, we assessed chemotaxis to *E. coli* plated on 0.2µm filter discs placed on the SoMo plate. The filter disc prevents bacteria from directly contacting the culture medium, but allows small molecules to diffuse through it. In this case, the chemotactic index was determined relative to a 0.2µm filter disc with no bacteria (Fig. 2B). *T. brucei* were attracted to *E. coli* grown on the 0.2µm filter, with a positive chemotactic index of + 0.42, indicating that attraction occurs in response to a factor smaller than 0.2µm that diffuses through the culture medium. It is important to note that the attractant may be a signal produced directly from the bacteria, or it may be a product of a chemical reaction between a factor produced by the bacteria and a substance present in the culture medium.

Because *T. brucei* are attracted to a diffusible cue emanating from *E. coli*, we next asked if this required dead or dying bacteria. When bacteria die, they often lyse, releasing intracellular metabolites into the environment, which can serve as nutrient sources for other microbes (29, 30), so we asked if *T. brucei* were attracted to products released from lysed *E. coli*. To determine the number of lysed bacterial cell equivalents to test, we determined the number of *E. coli* cells present 96 hpp (Fig. S2) because parasites show an attractive response to bacteria within 96 hpp (Fig. 1C). First, hypotonically lysed bacteria were tested to ask if the attractant might be a protein released from lysed bacteria, but no chemotactic effect was seen (Fig. 2A). Second, boiled *E. coli* cell lysates were tested. Boiling *E. coli* would denature proteins and inactivate heat labile compounds, but other potential metabolites would still be present; however, boiled lysates were repulsive to *T. brucei*. We also assessed the chemotactic index of dead but non-lysed *E. coli*, using formaldehyde-killed bacteria and found that *T. brucei* were repelled by formaldehyde-killed bacteria. Taken together, these results indicate that the attractant is diffusible through the culture medium, and actively growing bacteria are required for its production. Efforts to isolate the attractant have so far been unsuccessful.

### 3. Projections of parasites accelerate upon sensation of an attractant

The attraction of social *T. brucei* to *E. coli* presents an opportunity to investigate changes in *T. brucei* cell behavior underlying chemotaxis. In time-lapsed video analysis we noticed that projections appeared to speed up just before they made contact with the bacterial colony (Fig. 3A; Movies 1 and 2). To quantify this, we measured the distance each projection travelled between each frame of the time-lapsed video, either in the presence or absence of bacteria. A plot of distance traveled over time was then generated for each projection (Fig. 3B).

**Figure 3:**
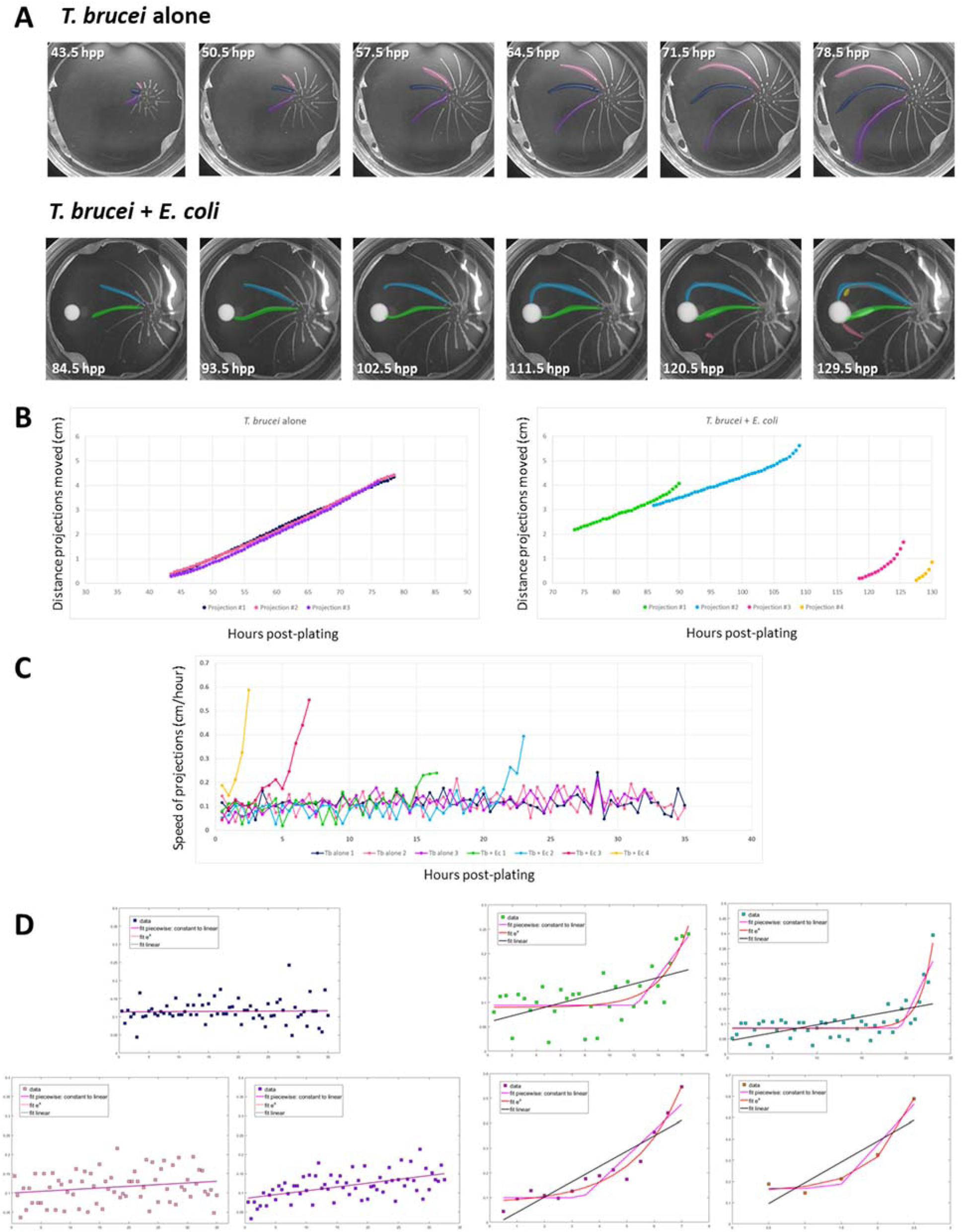
Projections of parasites accelerate upon sensation of attractant. A) Representative images of *T. brucei* engaging in SoMo alone (upper) (Movie 1) or with *E. coli* present (lower) (Movie 2). Projections are pseudo-colored and match the plot colors shown in panel B. Time-stamps are indicated in hours post-plating (hpp). B) The distance each projection moved was measured over time from the time-lapse videos shown in panel A. C) The speed of each projection is plotted over time with the colors of each projection corresponding to their respective colors shown in panels A and B. D) Non-linear regression models designed in MATLAB were used to model the best fit of the speed vs time data for projections in the presence of *E. coli*. The pink line represents the piece-wise function, the red line is for an exponential function, and the black line represents a linear function. The colors of the data points correspond to the respective pseudo-colors for each projection.

In the absence of bacteria, projections moved with mostly constant speed, as indicated by the constant slope of the line generated by the distance vs time analysis (Fig. 3B, left). Distance vs time measurements were fit to a linear or quadratic regression model (Table 1). All three projections fit the linear model very well (R^2^ = 0.99). Although the quadratic model also fit (R^2^ = 0.99), the constant in front of x^2^ value in each of the three equations was always very small, indicating that in the absence of bacteria, the projections do in fact move with a constant speed (∼0.1 cm/hr) (Table 1, Table S1). For projections moving in the presence of bacteria, the distance traveled versus time analysis indicated that the projections moved at a constant speed at first, as indicated by the constant slope at early time points. In the hours before impact with the bacteria however, the slope of the line continually increased, indicating that the projections of parasites were accelerating toward the bacteria (Fig. 3B, right).

**Table 1:** Equations of the regression models for T. brucei projections in the absence of bacteria: Distance vs Time. Equations for the best fit for both a linear regression and quadratic regression model were calculated for the indicated *T. brucei* projection in the absence of bacteria shown in Figure 3B. R^2^ values were calculated in Microsoft Excel for each regression analysis.

| Distance vs. Time graphs |  |  |
| --- | --- | --- |
| Projection | Linear regression:<br>$y = ax + b$ | Quadratic regression<br>$y = ax^2 + bx + c$ |
| *T. brucei* alone 1 | $y = 0.1172x - 4.8248$<br>$R^2 = 0.9996$ | $y = -2e-05x^2 + 0.1191x - 4.8819$<br>$R^2 = 0.9996$ |
| *T. brucei* alone 2 | $y = 0.1184x - 4.9054$<br>$R^2 = 0.9975$ | $y = 0.0006x^2 + 0.0441x - 2.7047$<br>$R^2 = 0.9997$ |
| *T. brucei* alone 3 | $y = 0.1209x - 5.1828$<br>$R^2 = 0.9965$ | $y = 0.0008x^2 + 0.027x - 2.4479$<br>$R^2 = 0.9995$ |

Because it is difficult to fit an equation to a line that changes from a constant slope to an increasing slope, we plotted the change in distance between each time point (i.e. the derivative), which represents the speed versus time (Fig. 3C). While there was noise in the speed versus time analysis for each projection measured, all projections in the presence of bacteria clearly increased their speed above the baseline speed of projections in the absence of bacteria (Fig. 3C, Fig. S3). For example, in the final 30 minutes before they impacted bacteria, projections reached speeds from 0.24 to 0.59 cm/hr (Fig. 3C).

To model the speed change of projections in the presence of bacteria, we used different non-linear regression models designed in MATLAB to analyze the speed versus time data (Fig. 3D, Table 2). Because projections appeared to move with a constant speed early and then accelerate before colliding with the bacteria, we first asked how well the data could be modeled by a piecewise function that began with a line with zero slope (i.e. constant speed), and then at an unknown time point (denoted by the term “k”) changed to a line with a constant and positive slope (i.e. increasing speed) (Table 2). We also asked how well the speed data fit an exponential equation, which would give an equation with an almost zero slope at early time points and then continuously change to an increasing slope. Finally, we also asked how well the data fit a linear equation, which would give an equation of a line with an unchanging slope. We found that while the piecewise function that modeled a zero slope to constant slope fit the speed data well (R^2^ = 0.522 – 0.929), the exponential regression provided a slightly better model for the data (R^2^ = 0.544 – 0.973) (Table 2). Linear regression did not fit the data as well as the other two models (Table 2, Table S1). These same models were applied to the speed of projections in the absence of bacteria, and most models gave equations of lines with close to zero slope, confirming that these projections move with a constant speed. These analyses indicate that in the presence of bacteria, projections move with a mostly constant speed, but in the hours before they reach the bacteria the group increases its speed dramatically. Analysis of additional time-lapse videos in the presence or absence of bacteria are consistent with these findings (Fig. S3, Table S1).

**Table 2:**
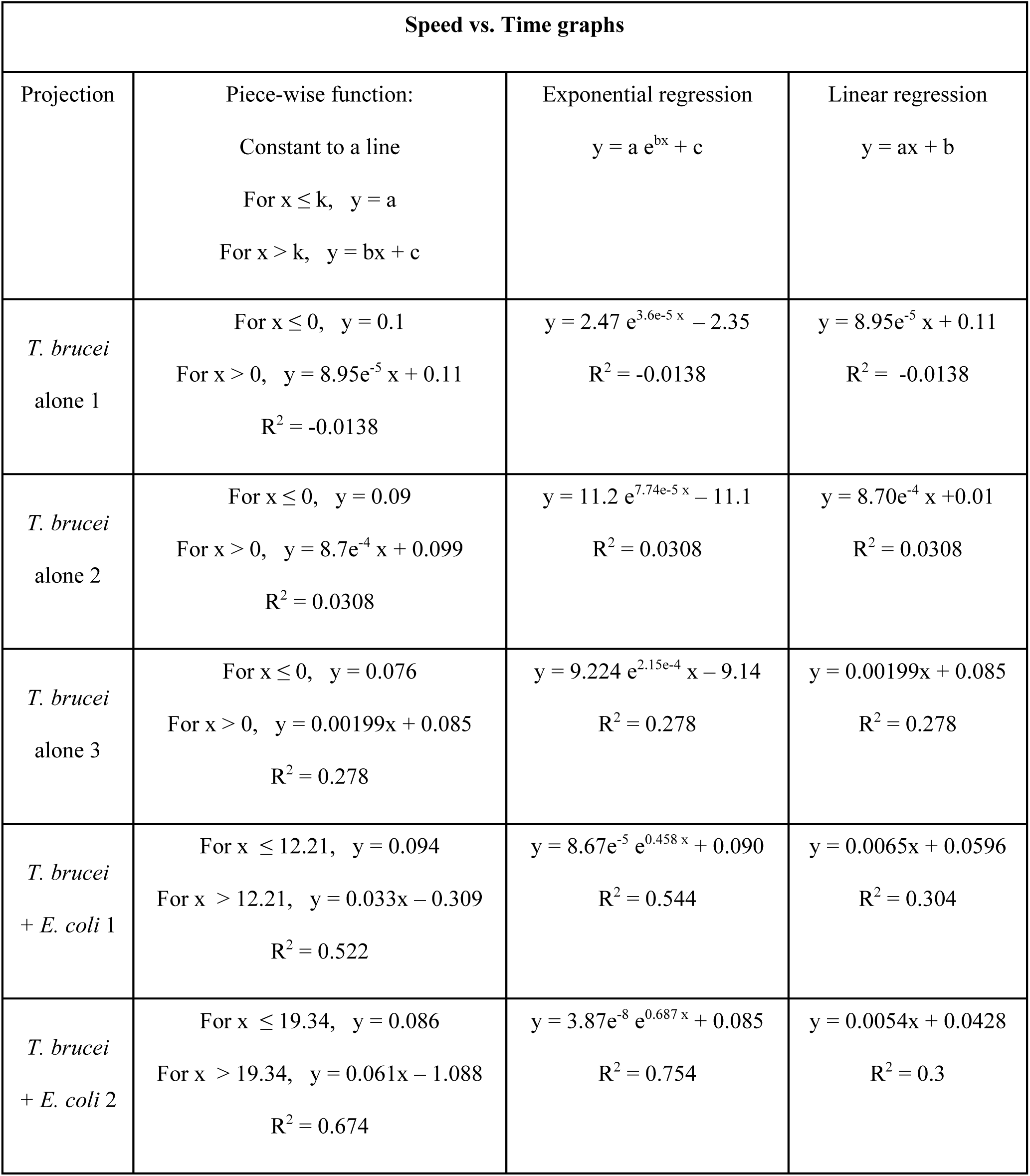
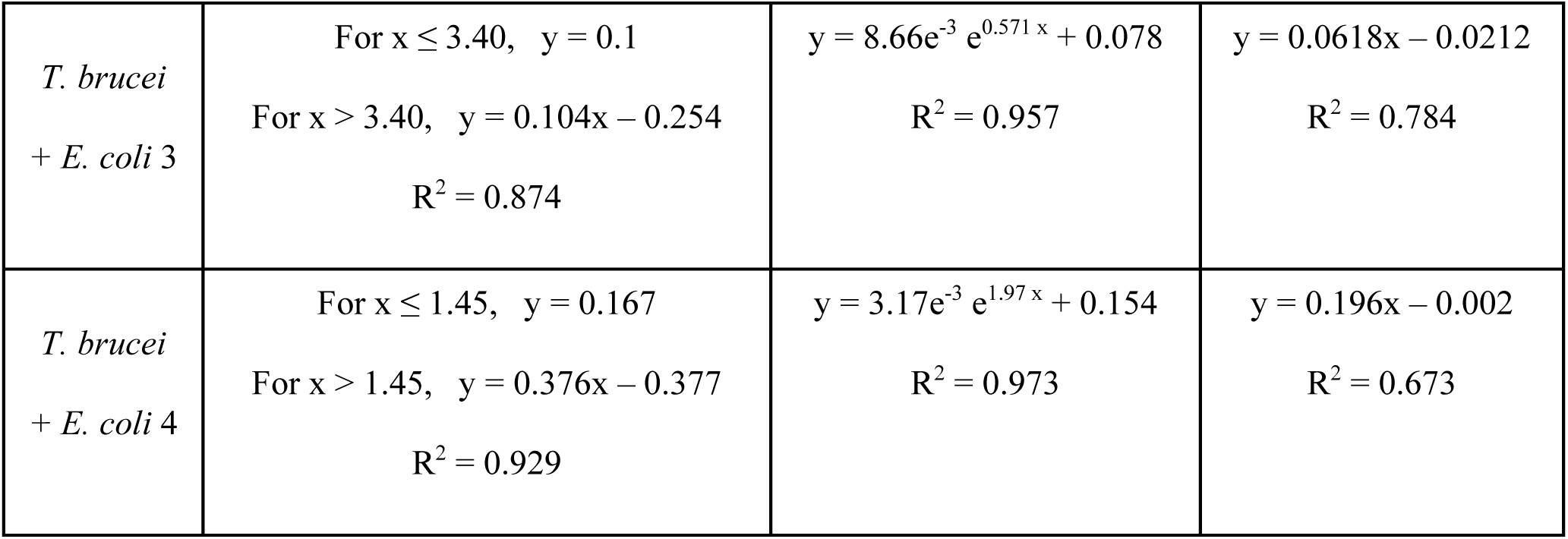
Equations of the regression models for T. brucei projections in the absence or presence of bacteria: Speed vs Time. Equations for the best fit for a non-linear fitting algorithm for a piece-wise, exponential, and linear function were calculated for the Speed vs Time data of *T. brucei* projections in the absence or presence of *E. coli*. R^2^ values were calculated by the non-linear fit model in MATLAB for each regression analysis.

| Speed vs. Time graphs |  |  |  |
| --- | --- | --- | --- |
| Projection | Piece-wise function:<br>Constant to a line<br>For $x \leq k$, $y = a$<br>For $x > k$, $y = bx + c$ | Exponential regression<br>$y = a e^{bx} + c$ | Linear regression<br>$y = ax + b$ |
| *T. brucei*<br>alone 1 | For $x \leq 0$, $y = 0.1$<br>For $x > 0$, $y = 8.95e^{-5}x + 0.11$<br>$R^2 = -0.0138$ | $y = 2.47 e^{3.6e^{-5}x} - 2.35$<br>$R^2 = -0.0138$ | $y = 8.95e^{-5}x + 0.11$<br>$R^2 = -0.0138$ |
| *T. brucei*<br>alone 2 | For $x \leq 0$, $y = 0.09$<br>For $x > 0$, $y = 8.7e^{-4}x + 0.099$<br>$R^2 = 0.0308$ | $y = 11.2 e^{7.74e^{-5}x} - 11.1$<br>$R^2 = 0.0308$ | $y = 8.70e^{-4}x + 0.01$<br>$R^2 = 0.0308$ |
| *T. brucei*<br>alone 3 | For $x \leq 0$, $y = 0.076$<br>For $x > 0$, $y = 0.00199x + 0.085$<br>$R^2 = 0.278$ | $y = 9.224 e^{2.15e^{-4}x} - 9.14$<br>$R^2 = 0.278$ | $y = 0.00199x + 0.085$<br>$R^2 = 0.278$ |
| *T. brucei*<br>+ *E. coli* 1 | For $x \leq 12.21$, $y = 0.094$<br>For $x > 12.21$, $y = 0.033x - 0.309$<br>$R^2 = 0.522$ | $y = 8.67e^{-5} e^{0.458x} + 0.090$<br>$R^2 = 0.544$ | $y = 0.0065x + 0.0596$<br>$R^2 = 0.304$ |
| *T. brucei*<br>+ *E. coli* 2 | For $x \leq 19.34$, $y = 0.086$<br>For $x > 19.34$, $y = 0.061x - 1.088$<br>$R^2 = 0.674$ | $y = 3.87e^{-8} e^{0.687x} + 0.085$<br>$R^2 = 0.754$ | $y = 0.0054x + 0.0428$<br>$R^2 = 0.3$ |
| *T. brucei*<br>+ *E. coli* 3 | For $x \leq 3.40$, $y = 0.1$<br>For $x > 3.40$, $y = 0.104x - 0.254$<br>$R^2 = 0.874$ | $y = 8.66e^{-3} e^{0.571x} + 0.078$<br>$R^2 = 0.957$ | $y = 0.0618x - 0.0212$<br>$R^2 = 0.784$ |
| *T. brucei*<br>+ *E. coli* 4 | For $x \leq 1.45$, $y = 0.167$<br>For $x > 1.45$, $y = 0.376x - 0.377$<br>$R^2 = 0.929$ | $y = 3.17e^{-3} e^{1.97x} + 0.154$<br>$R^2 = 0.973$ | $y = 0.196x - 0.002$<br>$R^2 = 0.673$ |

### 4. Individual cell motility within the group is constrained when sensing an attractant

To assess changes in individual cell behavior occurring in response to attractant, untagged wild type cells were mixed with 10% GFP-expressing cells. Through the use of a cell-tracking algorithm (31), the movements of individual GFP-tagged cells were assessed within the group. Individual cells at the tips of projections that were either not attracted to *E. coli* (n = 1368 cell tracks) or attracted to *E. coli* (n = 1403 cell tracks) were traced in 30-second videos (Fig. 4A). We assessed mean-squared displacement (MSD), which takes into account both the speed of cells and how far they move from their initial locations. We found that cells undergoing chemotaxis to *E. coli* had a lower MSD than cells in projections that were not undergoing chemotaxis (Fig. 4B). The lower MSD could mean that individual cells move more slowly in response to the attractant or that they alter how they move. To examine this further, we plotted the distribution of each cell’s curvilinear velocity versus straight-line velocity. We found that cells sensing an attractant had reduced straight-line velocity compared to cells not engaged in chemotaxis, suggesting that when an attractant is detected, parasites restrict their motion to smaller and more curving paths to remain near to the attractant (Fig. 4C). To quantify this change, we determined linearity for each cell, calculated as the ratio of straight-line velocity to curvilinear velocity. The mean linearity for cells not undergoing chemotaxis was 0.174, while the mean linearity for those undergoing chemotaxis was significantly decreased at 0.110 (two-tailed t-test, p < 0.0001). A similar paradigm, i.e. cells constraining their movements as they move closer to an attractant source, has been described for bacterial chemotaxis to K^+^ ions within a biofilm (32).

**Figure 4:**
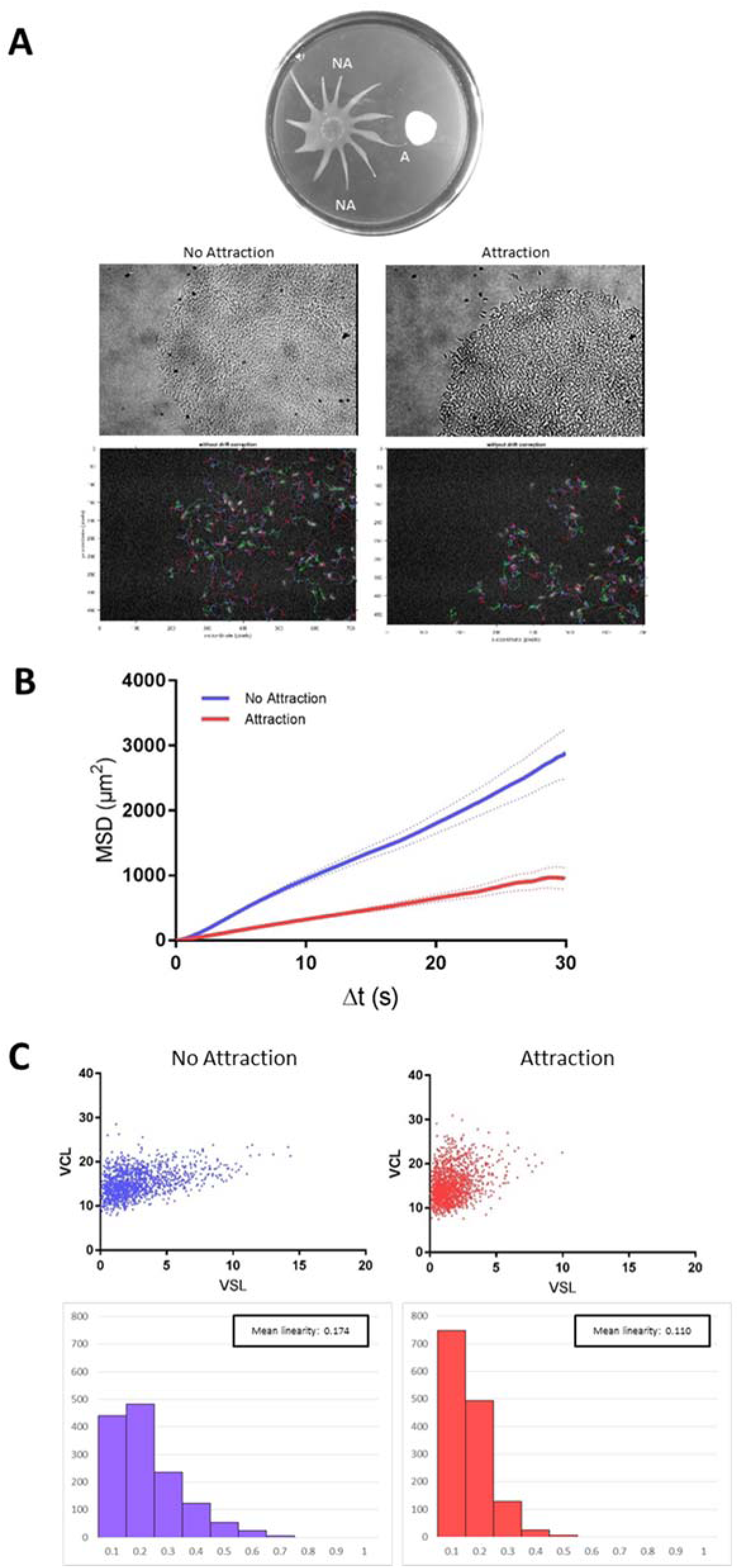
Individual cell motility within the group becomes more constrained in the presence of attractant. A) A representative SoMo plate is shown with a projection undergoing chemotaxis to *E. coli* (Attraction, “A”) and two projections not engaged in chemotaxis (No Attraction, “NA”). Representative phase contrast images of the tips of projections at 20x magnification are shown. Fluorescent images of the same tips of projections show GFP-tagged cells superimposed with their cell traces over a 30 second timeframe. B) Mean-squared displacement of individual cells at the tips of projections undergoing chemotaxis (Attraction) or not (No Attraction). Data acquired from 37 videos with 1368 total tracks (No Attraction), or 22 videos with 1403 total tracks (Attraction). C) Curvilinear versus straight line velocity plots with corresponding linearity plots are shown for Attraction and No Attraction conditions.

### 5. Group movement and single-cell movement are correlated

To ask if acceleration of projections toward *E. coli* correlates directly with constrained motility of individual cells, we monitored projection movement and individual cell motility in parallel as a function of time. SoMo assays were done with a mixture of 1% GFP-tagged cells mixed with untagged cells. 30-second movies of individual fluorescent cells at the tips of the projections were taken at the same time-points as photographs of projections over a two day time course. In a representative SoMo plate of *T. brucei* co-inoculated with *E. coli*, the distance projections moved was measured every 2 hours from 68.5 to 74 hours post-plating and from 90 to 96 hours post-plating, moments before the projection collided with the bacteria (Fig. 5A). As expected, projections increased their speed as they moved toward bacteria, accelerating from 0.0231 cm/hr to 0.1563 cm/hr (Figure 5C, Table 3). In contrast, in the absence of bacteria, the speed of the projection did not change substantially: 0.0315 cm/hr to 0.0502 cm/hr (Fig. 5D, Table 3). It should be noted that these speeds as slower overall than those seen in the time-lapse videos in Figure 3, which may be due to slight technical differences between the two assays. Over the time-course analysis of the projection tips became more curved (Fig. 5A and B, center), but this was observed regardless of whether bacteria were present (Fig. S2B), indicating it is likely a characteristic of advancing projections.

**Figure 5:**
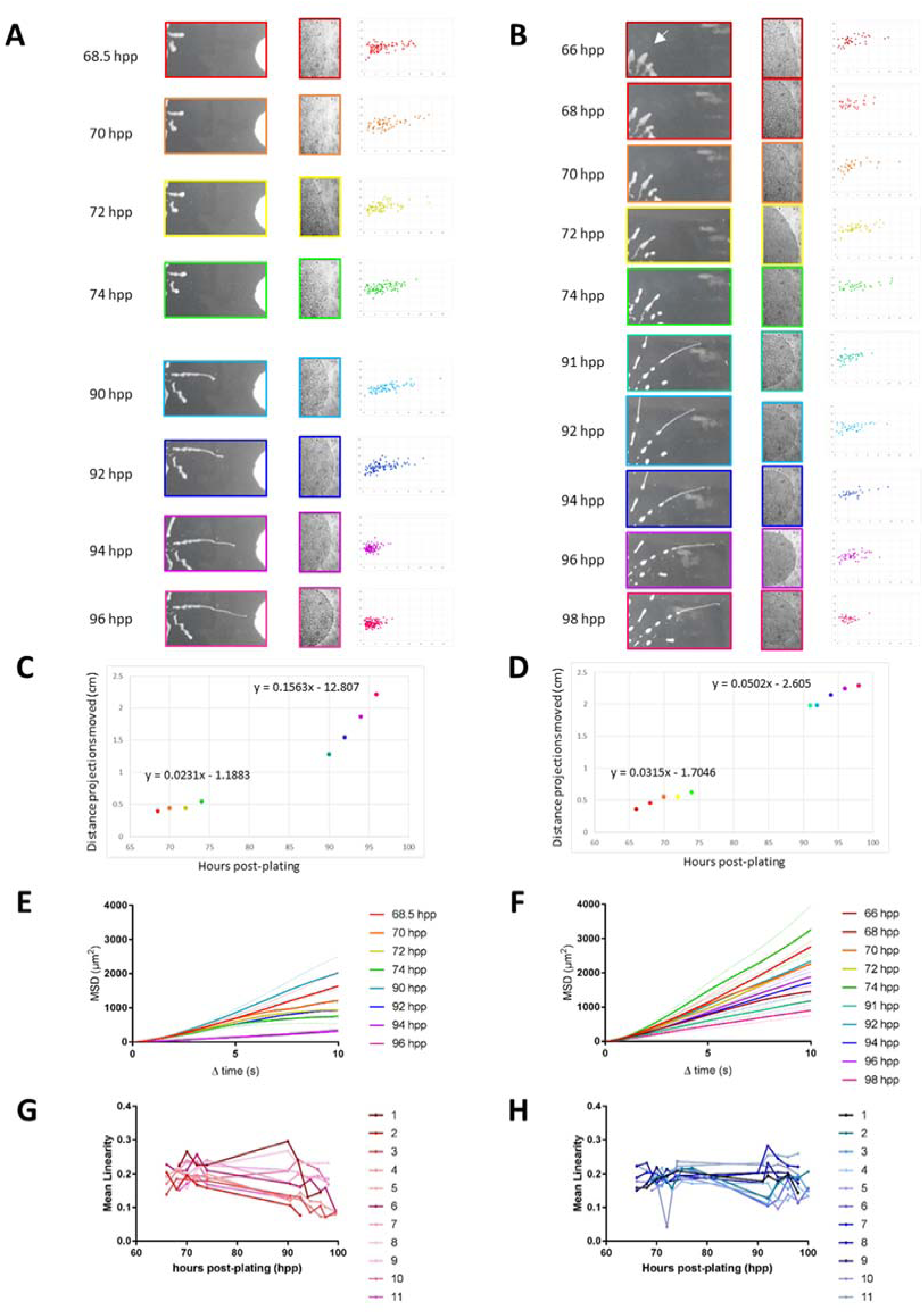
**Changes in group movement correlate with changes in single-cell movement during positive chemotaxis.** Speed of projections and MSD of individual cells within tips of these projections were examined over time in samples with or without *E. coli*. A) Sequential images of a projection moving toward *E. coli* (left column). At each time point, the tip of the projection is shown at 20x magnification (center column), and the curvilinear velocity vs straight-line velocity is shown for individual cells in the projection (right column). B) Projection of *T. brucei* alone analyzed as described in A. C and D) Distance from point of origin to the projection tip is plotted as a function of time for *T. brucei* projections moving in the presence (C) or absence (D) of *E. coli*. Equations for line-of-best fit using a linear regression are shown above the corresponding portions of each graph. E and F) Mean-squared displacement was determined for individual cells in the tip of each projection shown in A (C) and B (D). Line colors correspond to the colors used for time-points in A and B. G and H) Mean linearity of individual cells at the tips of projections was plotted over time for 11 projections each of *T. brucei* in the presence and absence of *E. coli*.

**Table 3:** Equations for the lines of best fit for T. brucei projections in the time-course analyses: Distance vs Time. Equations for the line of best fit for each section of the graphs in Figure 5C and D were determined using a linear regression analysis. R^2^ values were calculated in Microsoft Excel for each regression analysis.

| *E. coli* + *T. brucei* |  |
| --- | --- |
| 68.5 – 74 hpp | 90 – 96 hpp |
| Linear regression | Linear regression |
| $y = 0.0231x - 1.1883$<br>$R^2 = 0.8609$ | $y = 0.1563x - 12.807$<br>$R^2 = 0.9959$ |
| *T. brucei* alone |  |
| 66 – 74 hpp | 91 – 98 hpp |
| Linear regression | Linear regression |
| $y = 0.0315x - 1.7046$<br>$R^2 = 0.9245$ | $y = 0.0502x - 2.605$<br>$R^2 = 0.9572$ |

To ask how the movement of individual cells within the projections changed over time, we monitored the movement of individual cells in the same projections used for the speed analysis above. In the presence of *E. coli* parasite cells showed a decrease in MSD and exhibited increasingly constrained motility over time (Fig. 5A right and E), consistent with the results from Fig. 4B and C. While there was variation in the MSD of individual cells over time, a significant decrease in MSD was clear by 94 and 96 hpp (One-way ANOVA, p < 0.0001). Interestingly, the decrease in MSD seen in the presence of bacteria was not observed until the last two time-points before the projection impacted the bacteria (Fig. 5E, 94 and 96 hpp), even though the projection as a whole had already begun increasing its speed in the prior two time-points (Fig. 5C, 90 and 92 hpp). This finding suggests that, while changes in group and individual movement in the presence of bacteria are correlated, projection speed increases first, and individual cells constrain their motility later. We suspect there is some aspect of individual cell behavior that leads to the increased speed of the group as a whole, but so far, this has not been revealed in our current assays. In the absence of bacteria, *T. brucei* cells showed variation in MSD over time (Fig. 5F), which was reflected in variation of how constrained the movement of cells was over time (Fig. 5B right). However, no single time-point exhibited a significant decrease in MSD from all of the others (One-way ANOVA) (Fig. 5F), indicating that cells in the absence of bacteria were not changing how they moved within the projection, in contrast to how individual cells behave in the presence of bacteria.

When individual cells in multiple projections were monitored over time, this same trend held true (Fig. 5F and G). In the presence of bacteria, the mean linearity of cells in projections decreased over time (Fig. 5G), indicating that their movements were becoming more constrained as they moved toward the bacteria. In the absence of bacteria the mean linearity was variable and although some decrease occurred, the large drop seen during final time points in the presence of bacteria was not observed. (Fig. 5H). Taken together, these time-course experiments indicate that in response to bacteria, the speed of projections increases, and subsequently, individual cells within projections constrain their motion.

## Discussion

We have shown that socially behaving *T. brucei* engage in positive chemotaxis towards a diffusible cue produced by actively growing *E. coli*, and we have begun to elucidate the changes in group and individual cell behaviors that characterize this response. While chemotaxis has been well-studied in bacteria and other protozoans (1–3), to our knowledge, our results provide the first demonstration of positive chemotaxis in *T. brucei* and illustrate the capacity for interkingdom interactions between bacteria and protozoa.

In its tsetse fly host, *T. brucei* must undergo a series of specific migrations and differentiations as it moves from the fly midgut to the proventriculus, and finally to the salivary glands. It is therefore reasonable to anticipate that *T. brucei* may employ chemotaxis to direct movement in response to signals in specific fly tissues. Supporting this idea, prior work connected cAMP as a second messenger to control of parasite group movements *in vitro* (23, 24), and recent work demonstrated that *T. brucei* requires an intact cAMP signaling pathway in order to progress through tsetse fly tissues *in vivo* (8). The idea that chemotactic cues direct parasite progression through their insect vectors has been proposed for other kinetoplastid parasites, including *Leishmenia* spp and *Trypanosoma cruzi,* which have been shown to engage in chemotaxis *in vitro* (9, 11, 12). Our discovery of positive chemotaxis in *T. brucei* demonstrates that, in addition to moving away from external signals (19), these organisms can detect and move toward specific signals in their extracellular environment.

The ability to sense and respond to signals is also expected to be important for *T. brucei* within its mammalian hosts. Prior work has shown that in addition to the bloodstream and central nervous system, skin and adipose tissues represent important reservoirs contributing to pathogenesis and transmission (15–17). Mechanisms underlying *T. brucei* tropism to extravascular tissues remain to be determined, but positive chemotaxis could be involved. Study of the malaria parasite, *Plasmodium berghei*, in a mammalian host found that as parasites moved closer to blood vessels, their trajectories became more constrained (33), which mirrors the more constrained individual cell motility exhibited by *T. brucei* in response to attractant (Fig. 4C, 5A, 5G). Chemotaxis may also help *T. brucei* evade the immune system, analogous to what has been proposed for evasion of host neutrophils by *Leishmania* parasites (10). Motility is essential for *T. brucei* virulence in the mammalian host, perhaps allowing for quick changes in direction to avoid immune cells (31). Thus, the change in individual cell motility observed as parasites move toward an attractant (Fig. 4C, 5A, 5G) may be an important mechanism to infiltrate tissues and/or evade immune cells within the host.

The question arises as to why *T. brucei* exhibits chemotaxis toward bacteria. Notably, although *E. coli* is not typically present within the tsetse fly, the fly is home to three species of endosymbiotic Gram-negative bacteria: *Sodalis glossinidius*, *Wigglesworthia glossinidia*, and *Wolbachia* spp (34). While *Wolbachia* is restricted to the reproductive tract, *Wigglesworthia*, an obligate endosymbiont, is found intracellularly in bacteriocytes within a specialized structure called the bacteriome near the anterior midgut, and extracellularly in the milk glands (35, 36). *Sodalis* is found throughout the fly midgut and in a variety of other tissues (34). Evidence for functional interactions between *T. brucei* and bacterial endosymbionts of the tsetse comes from work demonstrating *T. brucei* reliance on bacterial products during fly transmission (34). For example, *Wigglesworthia* produces folate and phenylalanine, but *T. brucei* cannot. *T. brucei* does, however, encode transporters for these metabolites (37). Similarly, *T. brucei* encodes an incomplete threonine biosynthesis pathway (38). While the tsetse cannot provide homoserine that is necessary for *T. brucei* threonine biosynthesis (39), *Sodalis* can (40). A long-standing interaction between *T. brucei* and *Sodalis* is also evidenced by an example of horizontal gene transfer of the gene phospholipase A1 from *Sodalis* to *T. brucei* (41, 42). Of note, numerous studies of lab-reared and field-caught tsetse flies have demonstrated that the presence of *Sodalis* in the tsetse increases the likelihood of infection by *T. brucei* (43–46). Therefore, *T. brucei* likely interacts closely with *Sodalis* during the transmission cycle, and chemotaxis toward *Sodalis* would be advantageous. While *Sodalis* can be cultured *in vitro*, we were unable to culture them on *T. brucei* media, or vice versa, and thus the chemotactic index of *T. brucei* toward *Sodalis* could not be determined.

In addition to containing three endosymbiotic bacteria, the midgut of field-caught tsetse flies harbor a wide variety of bacterial species, with variation among different species of tsetse flies and geographic distribution (46). The most common genera of bacteria found in tsetse fly midguts included *Enterobacter*, *Enterococcus*, and *Acinetobacter* (46). While the three endosymbionts discussed above are all Gram-negative bacteria, other common genera found in the fly midgut encompass both Gram-negative and Gram-positive bacterial species. We did not detect attraction or repulsion toward the Gram-positive bacterium *B. subtilis* (data not shown). Future work will be needed to assess whether *T. brucei* shows chemotaxis to other examples of Gram-positive and Gram-negative bacteria.

The exact *in vivo* correlates of group movements observed in SoMo remain unclear. However, our findings, together with recent work (8, 20, 21, 23, 24, 47), clearly illustrate the value of SoMo for uncovering novel aspects of trypanosome biology that are relevant *in vivo*. The discovery here of BacSoMo, positive chemotaxis in *T. brucei*, and development of quantitative chemotaxis assays enabled us to begin investigating cellular mechanisms underlying directed movement in these pathogens. These systems enable studies at the scale of groups of cells and at the level of individual cells within groups. In bacteria, both scales of analysis have provided important insights. In *Pseudomonas* and *Myxococcus* for example, analyses of group chemotaxis led to the discovery of signaling molecules and systems that regulate bacterial social behavior and motility on a surface (48, 49). In cyanobacteria, studies of individual cell movements within a group revealed how movement of individuals can dictate the trajectory of the group during the phototactic response (50). Thus, our analyses of changes in both group and individual cell behavior in response to attractant set the stage for further study of the cellular and molecular mechanisms underlying these responses.

Looking forward, the quantitative chemotaxis assays reported here will serve as an important tool to enable dissection of cellular and molecular mechanisms used by trypanosomes to detect and respond to environmental signals. As a straightforward *in vitro* assay, screens for small molecules that inhibit or alter trypanosome sensory behavior without affecting parasite fitness could identify novel transmission blocking targets. Additionally, RNAi library screens for genes involved in sensory, signaling, or other chemotactic functions could be identified, further elucidating signaling systems required for parasite transmission and pathogenesis. Overall, the identification of positive chemotaxis in *T. brucei* will lead to deeper insights into how parasites sense and respond to cues in their changing host environments, facilitating the development of novel therapeutic and transmission blocking strategies.

## Materials and Methods

### Trypanosomes

*T. brucei brucei* 29-13 procyclic culture forms were used in this study (51). Parasites were cultured in SM media with 10% heat-inactivated fetal bovine serum (FBS) at 28°C and 5% CO_2_. GFP-tagged 29-13 cells were generated by transfection with Spe1-linearized pG-eGFP-Blast (gift of Isabel Roditi, University of Bern) as described previously (24). PDEB1 knockout cells were generated in the 29-13 background by two sequential rounds of homologous recombination using pTub plasmids conferring resistance to blasticidin and phleomycin (52). 452 bp upstream and 635 bp downstream of the PDEB1 coding sequence were used as regions of homology, the same regions used to independently create a PDEB1 KO cell line as in (8). Transfection of the knockout plasmids was performed as described previously (53). Primers used to amplify the PDEB1 regions of homology were described previously in (8), and are also listed here: Upstream FWD: atatGCGGCCGCTGCATTATGTTACTTGGGGGCA Upstream REV: atatCTCGAGGACGTAGTGTCCAACTGTGC Downstream FWD: atatGGATCCAGTCAGTTGACCGGTGGTAG Downstream REV: atatTCTAGACCGCCACAACTCCCTCTTAC

### Bacteria

*E. coli* strain DH5α with an ampicillin resistance plasmid were used for all experiments. *E. coli* from a glycerol stock were grown overnight in SM. 0.3µl of log-phase *E. coli* were inoculated on SoMo plates. Bacterial growth on SoMo plates was monitored over 4 days by washing colonies off independent SoMo plates twice per day and measuring the OD600 for 3 biological replicates.

### Social Motility assay

Plates for social motility assays were prepared based on (19). Briefly, a solution of 4% (wt/vol) SeaPlaque GTG agarose (Lonza) in MilliQ water was sterilized for 30 minutes at 250°C. Water that evaporated was replaced with sterile MilliQ water after heating, and the solution was then cooled to 70°C. SM made without antibiotics was pre-warmed to 42°C. The stock agarose solution was then diluted to 0.4% in the pre-warmed SM. The SM and agarose mixture was then mixed with ethanol (0.05% final solution) and methanol (0.05% final solution). 11.5 ml of the final mixture was poured in 100 mm by 15 mm petri dishes (Fisherbrand), which were allowed to dry with the lids off in a laminar flow hood for 1 hour.

Parasites in mid-log phase growth (1×10^6^ – 7×10^6^ cells/ml) were counted, harvested, and concentrated to 2 x 10^7^ cells/ml, and 5 µl of concentrated parasites were placed on the SoMo plate. Plates were allowed to incubate at room temperature for 10 minutes and were then wrapped in parafilm and placed at 28°C and 3% CO_2_. 24 hours post-plating, SoMo plates were moved to 28°C and 0% CO_2_.

### Chemotaxis assays

Chemotaxis assays were developed based on (26). SoMo plates, *T. brucei*, and *E. coli* were prepared as stated above. *T. brucei* was inoculated 1.5 cm from the center of the SoMo plate, and *E. coli* or the sample being tested was inoculated 2.0 cm from the center, opposite the *T. brucei*. At 120 hours post-plating, the number of projections that had entered a 2-cm diameter circle centered on the test sample location were counted. A chemotaxis index was calculated for each sample by comparing the average number of projections entering the circle for the sample condition to the average number of projections in control plates with *T. brucei* alone. Error bars represent standard error of the mean, and unpaired two-tailed t-tests were used to determine significance compared to the *T. brucei* alone control condition.

Sterilized 1-cm diameter Whatman filter discs were used for the Filter Paper condition. For the *E. coli* on lid condition, 9 ml of a 1.0% SM and agarose solution was plated on the lid of the SoMo plate and allowed to dry in the same manner as the SoMo plate. *E. coli* were then plated 2 cm from the center on the lid on the plate. The lid was placed on the plate such that the *E. coli* on the lid and *T. brucei* on the plate aligned on the same horizontal axis. For the 0.2 µm filter experiment, 0.2 µm GNWP Nylon membranes from Millipore were used. Because the 0.2 µm filters had a slightly larger diameter than the standard 2 cm diameter used in the other chemotaxis experiments, the number of projections that contacted the 0.2 µm filter were counted and compared in each condition.

For conditions in which *E. coli* cell lysate or formaldehyde-treated *E. coli* were tested, the number of bacterial cell equivalents that would have been present four days post-plating, as determined in our bacterial growth curve, were used. To generate hypotonically lysed *E. coli*, 1×10^11^ cells/ml from a log-phase overnight culture in LB were centrifuged at 8000 rpm for 10 minutes at 4°C. The supernatant was replaced with sterile MilliQ water. The sample was centrifuged again under the same conditions as above, and the supernatant was again replaced with water and lysozyme. The sample was then sonicated on ice 6 times for 10 seconds and then spun at 4000 rpm for 10 minutes to pellet the cell debris. The supernatant was filter-sterilized through a 0.2µm filter. For boiled *E. coli* lysate, 1×10^11^ cells/ml from a log-phase overnight culture in LB were centrifuged at 8000 rpm for 10 minutes at 4°C, and the supernatant was replaced with sterile MilliQ water. The sample was then boiled for 10 minutes. Finally, for formaldehyde-treated *E. coli*, 1×10^11^ cells/ml from a log-phase overnight culture in LB were centrifuged at 8000 rpm for 10 minutes at 4°C, washed in 1x phosphate buffered saline (PBS), and the PBS was replaced with 4% paraformaldehyde. Tubes with formaldehyde-treated *E. coli* were rotated at 4°C for 10 minutes, washed 3 times in 1x PBS, and spun down and re-suspended in the appropriate volume of 1x PBS.

### Time-lapse video analysis

SoMo assays were performed as described above. At 24 hours post-plating, SoMo plates were inverted and placed on a ring stand over a light box. Still photographs were taken every 30 minutes for 162 – 185 hours by a Brinno TLC200 Pro camera, positioned above the inverted plate. Camera settings were configured to compile still images into an .avi file with a play back speed of 10 frames/second. The .avi movies were converted into stacks using Fiji (Version 1.0) (54), and the segmented-line tool was used to measure the distance each projection moved over time.

### Non-linear regression analysis

A non-linear regression program was written in MATLAB to model the speed versus time data for the projections in the time-lapse videos. The “fit non-linear regression model” feature in MATLAB was used to create models for a piece-wise function that changed from a line with zero slope to a linear slope, an exponential function, and a linear function to find the best fit for the data.

### Individual Cell Motility analysis

For Figure 4, SoMo assays were performed as described above using a population of trypanosomes in which 10% GFP-tagged 29-13 cells were mixed with untagged cells and inoculated on the SoMo plate. Projections that had come near enough to *E. coli* to being moving toward it were placed in the “Attraction” category, and those that were not were placed in the “No Attraction” category. Tips of these projections were then imaged on a Zeiss Axiovert 200 M inverted microscope at 20x magnification under bright-field microscopy. Movies (30 seconds each) of fluorescent cells in these same projections were then captured at 30 frames per second with Adobe Premiere Elements 9 using 20x magnification under fluorescence microscopy. Fluorescent cells were tracked using a *T. brucei*-specific cell-tracking algorithm developed in MATLAB (31), and the resulting mean-squared displacement and curvilinear and straight-line velocities were calculated as described (55). Linearity is calculated as the ratio of straight-line velocity to curvilinear velocity. We only considered cells that were in focus for a minimum of 300 consecutive frames out of 900.

### Time-course analysis assessing projection movement simultaneously with individual cell motility

For time-course analyses in Figure 5, SoMo assays using 1% GFP-tagged cells mixed with untagged cells were performed as described above. At each time point indicated, SoMo plates were photographed using a Fujifilm FinePix JZ250 digital camera, tips of projections were imaged on a Zeiss Axiovert 200 M inverted microscope as described above, and a 30-second video of GFP-expressing cells was captured as described above for each projection. This analysis was done for 11 projections each for *T. brucei* alone and *T. brucei* + *E. coli*.

### Projection Curvature calculation

Bright-field images at 20x magnification were acquired for the tips of projections. A straight line of 3-inch standard length was used to measure the angle of curvature of the tip of projections. One end of the standard was placed tangent to the peak of the projection tip, and a straight line was drawn at the other end, perpendicular to the standard until it intersected with the projection. The interior angle was then calculated and assigned as the angle of curvature (Fig. S2C).

### Data Availability

All source data used to generate the figures in this manuscript are included in Table S2. Source code for the MATLAB modeling used to create Figure 3D, Figure S3D, Table 2, and the Speed vs Time table in Table S1 can be found at https://gist.github.com/614a9d210af934b2cedcc3a76f1b66f1.git. Data in this paper will be made fully available and without restriction.

## Acknowledgments

We thank Hunter Bennet for technical assistance with initial studies, and Michael Albanese for help developing non-linear regression models utilizing MATLAB. For their numerous helpful suggestions and discussions, we thank Drs. Wenyuan Shi and Marvin Whiteley as well as Dr. Beth Lazazzera, whom additionally provided the *B. subtilis* flagellar mutant. Members of the Hill laboratory are thanked for discussions and comments on the manuscript. K.L.H. NIH grant AI052348. S.F.D. Ruth L. Kirschstein National Research Service Award GM007185 and Ruth L. Kirschstein National Research Service Award AI007323. E.A.S. NIH-USPHS-NRSA GM07104. M.A.L. NIH-NRSA F31AI085961.

## Supplemental Material Legends

**Figure S1: *T. brucei* projections thicken prior to branching.**

Stills from a time-lapse video of *T. brucei* engaging in SoMo (Movie 3). Arrow points to the projection that thickens before branching. Time-stamps are indicated in hours post-plating (hpp).

**Figure S2: Growth analysis of *E. coli* colonies on SoMo plates, and the curvature of the tip of *T. brucei* projections increases in the presence and absence of *E. coli***

A) Growth of *E. coli* colonies was measured on SM supplemented agarose plates over the course of five days.

B) The angle of curvature of the tip of the projections in panels A and B of Figure 5 are plotted over time (hours post-plating).

C) A schematic of how the angle of curvature was determined in panel B is shown.

**Figure S3: Velocity analysis of projections from additional time-lapse videos (Movies 3 and 4) in both the presence and absence of bacteria.**

A) Representative images of *T. brucei* engaging in SoMo alone (upper) (Movie 3) or *T. brucei* engaging in SoMo with *E. coli* present (lower) (Movie 4). Projections are pseudo-colored and match the colors shown in panel B. Time-stamps are indicated in hours post-plating (hpp).

B) The distance each projection moved was measured over time from the time-lapse videos shown in panel A.

C) The speed of each projection is plotted over time with the colors of each projection corresponding to their respective colors shown in panels A and B.

D) Non-linear regression models designed in MATLAB were used to model the best fit of the speed vs time data for projections in both the presence and absence of *E. coli*. The pink line represents the piece-wise function, the red line is for an exponential function, and the black line represents a linear function. The colors of the data points correspond to the respective pseudo-colors for each projection.

**Table S1: Equations of the regression models for T. brucei projections from Movie 3: Distance vs Time, and for Movies 3 and 4: Speed vs Time.**

Equations for the best fit for both a linear regression and quadratic regression model were calculated for the indicated *T. brucei* projection in the absence of bacteria shown in Figure S3B. R^2^ values were calculated in Microsoft Excel for each regression analysis. Additionally, equations for the best fit for a non-linear fitting algorithm for a piece-wise, exponential, and linear function were calculated for the Speed vs Time data of *T. brucei* projections in the presence and absence of *E. coli*. R^2^ values were calculated by the non-linear fit model in MATLAB for each regression analysis.

**Table S2: Source data for all experiments.**

**Movie 1: Time-lapse video of *T. brucei* engaging in SoMo.**

Still images were taken every 30 minutes for 162 hours beginning 24 hours post-plating. Still images playback at 10 frames per second.

**Movie 2: Time-lapse video of *T. brucei* engaging in positive chemotaxis toward *E. coli*.**

Still images were taken every 30 minutes for 185 hours beginning 24 hours post-plating using a Brinno TLC200 Pro time-lapse camera. Still images playback at 10 frames per second.

**Movie 3: Time-lapse video of *T. brucei* engaging in SoMo.**

Still images were taken every 30 minutes for 161.5 hours beginning 24 hours post-plating using a Brinno TLC200 Pro time-lapse camera. Still images playback at 10 frames per second.

**Movie 4: Time-lapse video of *T. brucei* engaging in positive chemotaxis toward *E. coli*.**

Still images were taken every 30 minutes for 183.5 hours beginning 24 hours post-plating using a Brinno TLC200 Pro time-lapse camera. Still images playback at 10 frames per second.

**Movie 5: Live video of *T. brucei* cells at the tip of a projection upon initial impact with an *E. coli* colony.**

Cells were imaged at 20x magnification, and video was recorded and played back at 30 fps.

**Movie 6: Time-lapse video of a projection impacting an *E. coli* colony**

Still images were taken every 1 second at 5x magnification using AxioVision 4.7.2 (12–2008) and played back at 5 frames per second.

